# Permissive and instructive *Hox* codes govern limb positioning

**DOI:** 10.1101/2024.07.15.603511

**Authors:** Yajun Wang, Maik Hintze, Jinbao Wang, Hengxun Tao, Patrick Petzsch, Karl Köhrer, Longfei Cheng, Peng Zhou, Jianlin Wang, Zhaofu Liao, Xufeng Qi, Dongqing Cai, Thomas Bartolomaeus, Karl Schilling, Joerg Wilting, Stefanie Kuerten, Georgy Koentges, Ketan Patel, Qin Pu, Ruijin Huang

## Abstract

The positioning of limbs along the anterior-posterior axis varies widely across vertebrates. The mechanisms controlling this feature remain to be fully understood. For over 30 years, it has been speculated that *Hox* genes play a key role in this process but evidence supporting this hypothesis has been largely indirect. In this study, we employed loss- and gain-of-function *Hox* gene variants in chick embryos to address this issue. Using this approach, we found that *Hox4/5* genes are necessary but insufficient for forelimb formation. Within the *Hox4/5* expression domain, *Hox6/7* genes are sufficient for reprogramming of neck lateral plate mesoderm to form an ectopic limb bud, thereby inducing forelimb formation anterior to the normal limb field. Our findings demonstrate that the forelimb program depends on the combinatorial actions of these *Hox* genes. We propose that during the evolutionary emergence of the neck, *Hox4/5* provide permissive cues for forelimb formation throughout the neck region, while the final position of the forelimb is determined by the instructive cues of *Hox6/7* in the lateral plate mesoderm.

**Impact statement:** Elucidation of the *Hox* code defining forelimb positioning provides novel insights in lateral plate mesoderm patterning and the integration of vertebrate column structure and limb positioning.

## Introduction

The spatial development of vertebrate tissues is regulated by *Homeobox* (*Hox*) genes (Duboule, 2022; Iimura & Pourquie, 2006; Zakany & Duboule, 2007). A huge literature evidences that *Hox* genes determine the development and patterning of the vertebrate axial skeleton (reviewed in Burke, 2000). Mutations in *Hox* genes can lead to homeotic transformations, where one type of vertebra is transformed into another (Böhmer, 2017). Vertebrate limbs emerge at specific axial levels along the anterior-posterior (AP) axis, with precise positioning varying significantly across species (Burke et al., 1995). These characteristics make limb positioning a valuable experimental model for studying the mechanisms regulating positional information (Zakany & Duboule, 2007). Despite variable numbers of cervical vertebrae between species, the pectoral fin or forelimb is always located at the cervical-thoracic boundary. The mechanisms underpinning the positioning of vertebrate forelimbs remain to be fully elucidated.

While *Hox* gene misexpression causes substantial alterations in vertebrae identity (Garcia-Gasca & Spyropoulos, 2000; Horan et al., 1995; Jeannotte et al., 1993; Ramfrez-Solisn et al., 1993), only minor changes in limb development have been observed in *Hox* gene mutants (Rancourt et al., 1995). It is therefore unclear whether, for example, the abnormal limb that develops in *Hoxb5* mutants represents a true shift in the limb field or rather a shoulder girdle defect causing the forelimb to appear “shrugged” anteriorly. In fact, whereas knockdown of the complete paralogous group Hox5 genes results in changes in limb patterning, it does not result in a positional shift of the forelimb (Xu et al., 2013).

Moreover, interpretation of the effects of *Hox* genes on limb positioning in global knockouts is fraught by the fact that this not only affects lateral plate mesoderm patterning, but invariably also vertebrae-forming mesoderm and vertebrate identity. Yet normal vertebrate identity is required as a reference for defining limb positions. Ideally, limb positioning should be investigated by limiting the manipulation of *Hox* expression to the limb-forming mesoderm, without altering vertebral positional identity.

The initiation of the forelimb program is marked by *Tbx5* expression in the LPM, which is functionally required for pectoral fin formation in zebrafish and forelimb formation in chicken and mice (Hasson, Del Buono, & Logan, 2007; Rallis et al., 2003; Takeuchi et al., 2003). However, the forelimb-forming potential is present in mesodermal cells at the cervico-thoracic transitional zone long before the activation of *Tbx5* expression (Chaube, 1959; Moreau et al., 2019). This has led to the notion that cells first acquire positional identity through the expression of *Hox* genes, followed by a developmental program guided by their positional history (Duboule, 2022; Iimura, 2006; Zakany and Duboule, 2007).

The positional identity of future limb forming cells of the LPM is coded by the nested and combinatorial expression of *Hox* genes (Duboule & Dollé, 1989; Kessel & Gruss, 1991). Only a few studies have investigated how this *Hox* code translates to *Tbx5* expression in the prospective forelimb region and thus regulates forelimb positioning (Moreau et al., 2019). During gastrulation, the collinear activation of *Hox* genes begins in the epiblast, conferring anterior-posterior identity to the paraxial mesoderm (Duboule, 2022; Iimura & Pourquie, 2006). A similar mechanism regulates the anteroposterior patterning of the LPM, which gives rise to limbs. For instance, *Hoxb4*-expressing cells emigrating from the posterior part of the primitive streak form the LPM in the neck. Subsequently, *Hoxb4* activates *Tbx5* expression within this LPM domain.

The limb positioning is thus regulated by *Hox* genes in two phases (Minguillon et al., 2012; Moreau et al., 2019; Nishimoto et al., 2014). During the first phase, *Hox*-regulated gastrulation movements establish the forelimb, interlimb and hindlimb domains in the LPM. In the second phase, a *Hox* code regulates *Tbx5* activation in the forelimb-forming LPM (Minguillon et al., 2012; Moreau et al., 2019; Nishimoto et al., 2014). The forelimb-forming *Hox* code is considered to be constituted by both repressing and enhancing *Hox* genes: Caudal *Hox* genes, including *Hox9*, suppress and thus limit *Tbx5* expression, whereas rostrally expressed *Hox* genes activate *Tbx5* expression (Minguillon et al., 2012; Nishimoto et al., 2014). To date, *HoxPG4* and *PG5* genes are considered as activators of *Tbx5* (Minguillon et al., 2012; Nishimoto et al., 2014). The function of *PG6* and *PG7* genes (Becker et al., 1996; Becker, Jiang, et al., 1996), which are also prominently expressed in the forelimb region, has so far not been analysed.

Here, we aimed to investigate which *Hox* genes act to position the anterior limb in chicks. We present evidence that wing position is controlled by a permissive signal governed by *HoxPG4/5* which demarcates a territory where it can form. However, an addition instructive cue mediated by *HoxPG6/7* genes within the permissive region is required for forelimb formation. Our study is the first to show that neck LPM can be re-specified to form limb.

## Results

### *HoxPG4–7* are required for the forelimb formation

The expression domain of *HoxPG6/7*, like that of *HoxPG4/5*, overlaps with the forelimb field, suggesting they might activate *Tbx5* expression. To untangle the roles of individual members of the *HoxPG4/5/6/7*, we performed loss-of-function experiments in chick embryos. We focused on the A-cluster of *HoxPG4/5/6/7*, using specifically generated dominant-negative (DN) forms to suppress the signalling function of each target *Hox* gene. The DN variants lack the C-terminal portion of the homeodomain, rendering them incapable of binding to the target DNA while preserving their function of binding transcriptional specific co-factors (Denans et al., 2015; Gehring et al., 1990).The specificity and effectiveness of this dominant-negative strategy have been further validated in a recent study showing that expression of a Hoxb4 DN construct led to a reduction in the Tbx5 expression domain during limb induction (Moreau et al., 2018), consistent with a specific loss of Hoxb4 function. Plasmids expressing dominant-negative *Hoxa4, a5, a6* or *a7* were electroporated into the dorsal layer of LPM in the prospective wing field, from which the wing mesoderm originates in Hamburger-Hamilton stage (HH) 12 chick embryos (Hamburger & Hamilton, 1951) (Fig. 1a, b). After 8 – 10 h, embryos reached HH14 when expression from the transfected DN-constructs was detectable in the wing field of the transfected (right) side signified by Enhanced Green Fluorescent Protein (EGFP) expression also encoded by these plasmids (Fig. 1c).

**Fig. 1.**
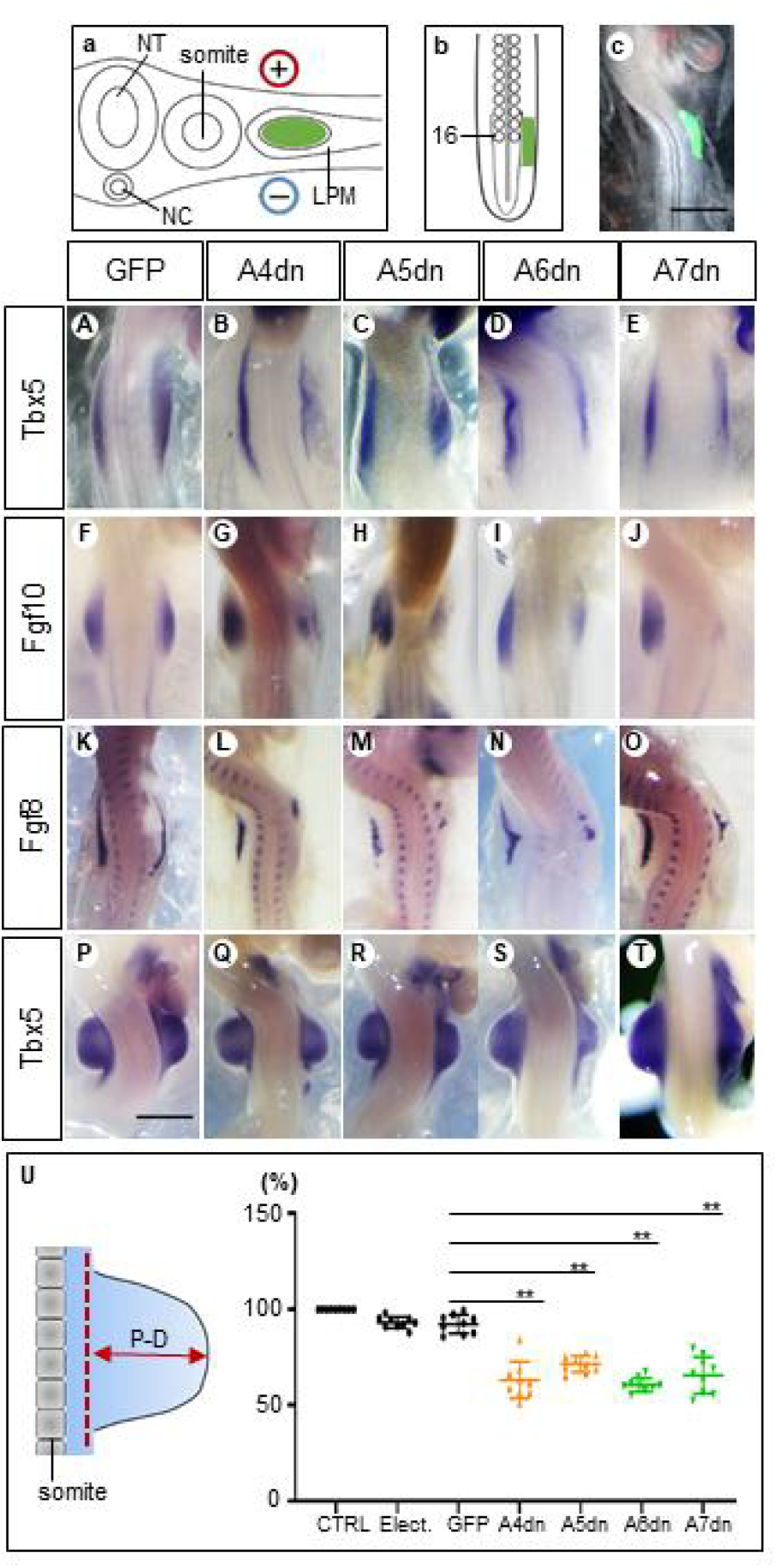
Hoxa4/a5/a6/a7 genes are necessary for wing bud formation. Schemes showing the electroporation in transverse section (**a**) and in the dorsal view (**b**). The somite 16 is marked (**b**). Successful transfection of plasmids as verified by EGFP expression(**c**). The *dn Hox* genes downregulated the expression of Tbx5 (**B**-**E**), Fgf10 (**G**-**J**), and Fgf8 (**L**-**O**) and inhibited wing bud formation at the ipsilateral (right) side (**Q**-**T**). **A**-**E**: HH14; **F**-**O**: HH18-19; **P**-**T**: HH22; scale bars in **c** (for **c, A**-**O**) and in **P** (for **P**-**T**): 500μm. The proximodistal (P-D) distance (left in **U**) of wing buds is significantly reduced in *Hox dn*-expressing wing buds compared to EGFP electroporated wing buds (right in **U**). The scheme on the left-hand side shows how measurements were made. Red dotted line: baseline of the wing bud; CTRL: normal control wing buds without any operation; Elect.: wing buds after electroporation without constructs; GFP: wing buds after electroporation with EGFP-expressing constructs; A4dn, A5dn, A6dn, A7dn: wing buds after electroporation with *dn* Hoxa4/5/6/7 expressing constructs, respectively. Each dot represents one embryo; error bars represent mean ±SEM. **p < 0.01.

*Tbx5* as the first gene indicating activation of the forelimb-forming program starts to be expressed in the forelimb field of normal chick embryos at HH13 (http://geisha.arizona.edu). Therefore, we analysed *Tbx5* expression following inhibition of *Hoxa4/5/6/7* at HH14. Expression of *Tbx5* in the wing field transfected with DN plasmids for any of these genes was consistently lower than in the contralateral (control) side (Fig. 1B-E, Table 1). The down-regulation of *Tbx5* expression by all four dominant-negative forms of *Hoxa4/5/6/7* shows a previously unknown requirement of *PG6* and *PG7 Hox* genes for the activation of *Tbx5* during forelimb induction, and confirms the previously reported *Tbx5* activating effects of *PG4* and *PG5 Hox* genes (Nishimoto et al., 2014)

**Table 1.** Dominant negative expression of Hoxa4/a5/a6/a7 down-regulated gene expression. The numbers of embryos with an unambiguous effect and the total number of embryos analysed are given (effect/total number analysed).

|  | Tbx5 |  |  | Fgf10 |  |  | Fgf8 |  |  |
| --- | --- | --- | --- | --- | --- | --- | --- | --- | --- |
|  | Expres<br>sion<br>level<br>decrea<br>sed | Express<br>ion<br>domain<br>shorten<br>ed | Both<br>down-<br>regulati<br>on | Expres<br>sion<br>level<br>decrea<br>sed | Expres<br>sion<br>domain<br>shorten<br>ed | Both<br>down-<br>regulati<br>on | Expres<br>sion<br>level<br>decrea<br>sed | Expres<br>sion<br>domain<br>shorten<br>ed | Both<br>down-<br>regulati<br>on |
| A4dn | 15/17 | 12/17 | 10/17 | 10/11 | 9/10 | 9/10 | 10/10 | 6/10 | 6/10 |
| A5dn | 6/6 | 5/6 | 5/6 | 5/6 | 5/6 | 5/6 | 4/6 | 4/6 | 4/6 |
| A6dn | 6/6 | 6/6 | 6/6 | 6/6 | 5/6 | 5/6 | 5/5 | 5/5 | 5/5 |
| A7dn | 5/8 | 4/8 | 5/8 | 6/8 | 6/8 | 6/8 | 6/6 | 5/6 | 5/6 |

*Tbx5* is required for the activation of *Fgf10* in the mesoderm (Cohn et al., 1995; Min et al., 1998; Sekine et al., 1999; Young et al., 2019). *Fgf10* subsequently induces *Fgf8* expression in the overlying ectoderm to initiate forelimb outgrowth (Barrow et al., 2003). The two genes form a positive feedback loop to ensure formation of the apical ectodermal ridge (AER), which ultimately regulates sustainable outgrowth and patterning (Crossley et al., 1996; Min et al., 1998). We analysed their expression at HH18-19. Dominant-negative inhibition of any of the *Hoxa4/5/6/7* genes reduced the expression levels and domains of *Fgf10* (Fig. 1G-J; Table 1) and *Fgf8* (Fig. 1L-O; Table 1). The consequence of these manipulations on the outgrowth of the wing bud was analysed at HH22, when the wing bud develops a nearly square shape. After inhibition of HOX proteins, the form of the target wing buds was altered and their size was decreased. In some cases, the anteroposterior extent of the wing bud was also remarkably reduced (Fig. 1Q-T). To quantify the effect of Hox inhibition on wing bud development, we measured the proximal-distal (P-D) elevation of the electroporated wing bud above the trunk lateral surface, compared to the contralateral non-electroporated control wing bud (Fig. 1U).

Electroporation of plasmid-free solution and an EGFP-encoding plasmid caused only minimal reduction compared to their contralateral wing bud, indicating low developmental toxicity of the procedure of electroporation itself (Fig. 1U).

Interference with the action of the representative A-cluster *Hox* genes indicate that *Hox* genes from all four paralogous groups (*PG4, PG5, PG6* and *PG7*) impinge on the forelimb program and should be considered part of the activating *Hox* code for forelimb development. Overall, the effect of each *Hox* gene is limited, suggesting they act in a combinatorial, and possibly redundant fashion.

### *Hox6/7* but not *Hox4/5* are sufficient to reprogram neck to wing mesoderm

We next investigated the role of *HoxPG6/7* during forelimb fate determination. We hypothesized that if *HoxPG6/7* are (an) integral and necessary part(s) of the forelimb *Hox* code, their ectopic expression in a non-limb region, similar to the limb-inducing activity of FGFs (Cohn et al., 1995), should induce forelimb formation. In the present study, the neck was chosen as the non-limb region.

When A-cluster genes were electroporated at HH11–12 into the dorsal LPM at the level of somites 10–14 (anterior to the wing field) (Fig. 2a), strong expression could be verified anterior to the cognate wing field by in situ hybridization (ISH) 12h after electroporation (Fig. 2b-e), indicating successful expression of *Hox* gene constructs.

**Fig. 2.**
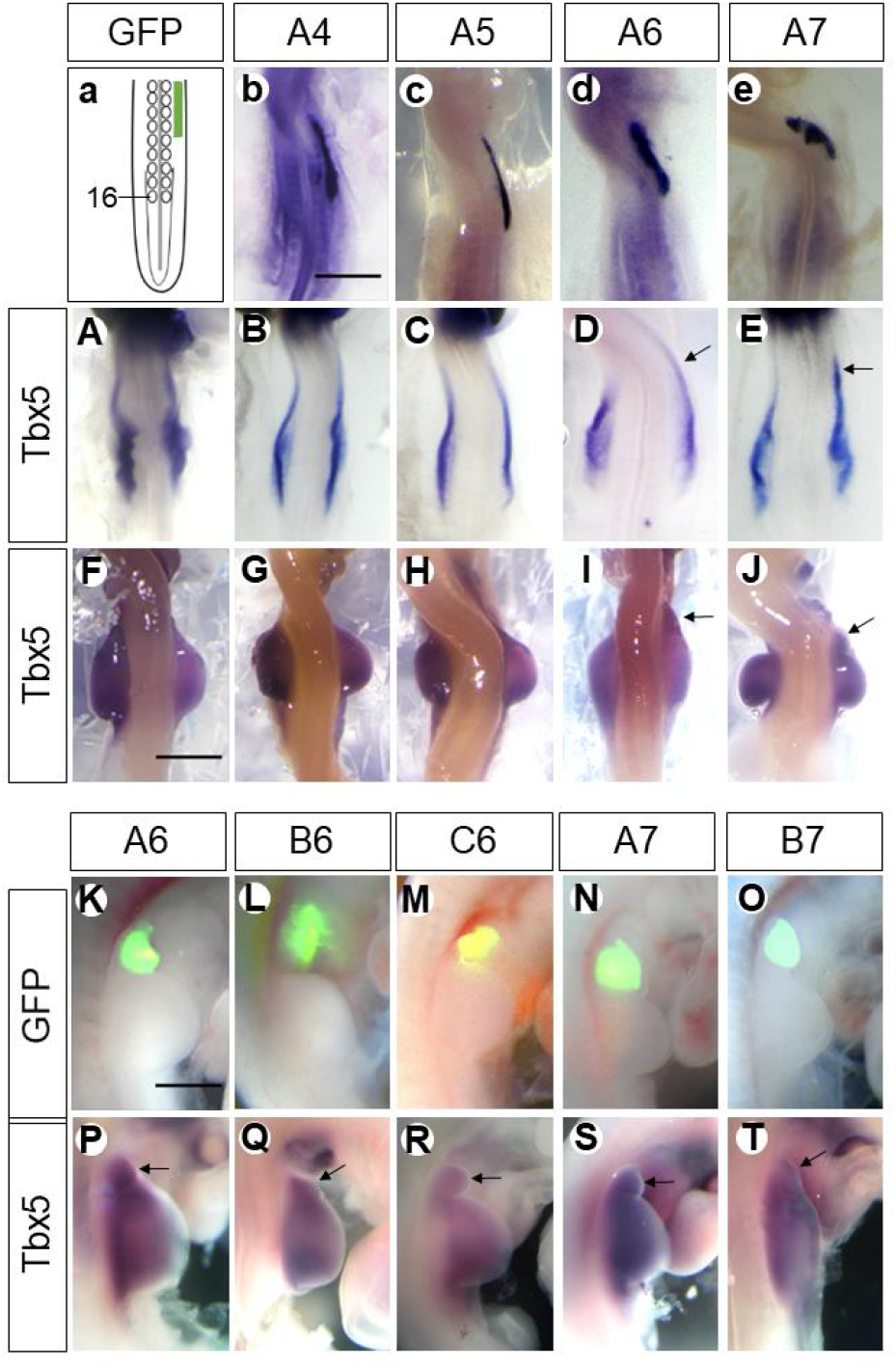
*Hoxa6/a7* but not *Hoxa4/a5* are sufficient to induce a neck wing bud. Scheme showing electroporation of the neck region in the dorsal view (**a**). The somite 16 is marked. The expression domain of electroporated constructs is marked by a green bar. Expression of *Hoxa4*(**b**), *Hoxa5*(**c**), *Hoxa6*(**d**), *Hoxa7*(**e**) in the LPM anterior to the wing field after electroporation with the respective plasmids as documented by in situ hybridization. Whereas ectopic cervical expression of *Hoxa6/a7* induced the anterior expression (indicated by arrows) of *Tbx5* (**D, E, I, J**), overexpression of *Hoxa4/a5* did not induce anterior expression of it (**B, C, G, H**). Also, only Hoxa6 and Hoxa7, but not a4 or a5 resulted in the anterior extension of the wing bud (arrows in **I**-**J**). The ectopic wing buds (fused with or separated from the endogenous one) induced by *HoxPG6-7* are indicated by GFP fluorescence (**K**-**O**) and in situ hybridization for *Tbx5* (arrows in **P**-**T**). **b**-**e** and **A-E**: HH14; **F-T**: HH22; scale bars in **b** (for **b-e** and **A**-**E)**, in **F** (for **F**-**J**) and in **K** (for **K**-**T)**: 500μm. Arrows indicate induced wing buds (**P-T**).

The anterior expression domain of *HoxPG6/7* overlaps with the forelimb field but does not extend into the neck region. Electroporation of constructs expressing *Hoxa6/7* into the neck mesoderm caused their ectopic expression anterior to the forelimb field (Fig. 2d, e). This induced ectopic expression of *Tbx5* in this region anterior to the cognate wing field (Fig. 2D, E). By 48h re-incubation, a bulge appeared in the neck region transfected with *Hoxa6/7*. This bulge expressed the forelimb master gene *Tbx5*, and expression strength was similar to that of the natural wing bud (Fig. 2I, J). Hence, it can be considered as an ectopic wing bud in the neck.

In contrast to ectopic expression of *Hoxa6/7* in the neck region, overexpression of *Hoxa4/5* (Fig. 2b, c) by electroporating this region with *Hoxa4/5* coding plasmids did not extend *Tbx5* expression anteriorly (Fig. 2B, C), indicating that no wing-forming mesoderm was ectopically induced in the neck by *Hoxa4*/*5* overexpression. Consequently, no structure emerged from the neck anterior to the endogenous wing bud after 48 h of re-incubation (Fig. 2G, H). These results demonstrate that *Hoxa4* and *Hoxa5* are insufficient, whereas *Hoxa6* and *Hoxa7* are sufficient to specify wing mesoderm in the neck region.

To ascertain whether other members of *HoxPG6/7* share the forelimb-inducing activity of the A-cluster genes, plasmids encoding full-length *Hoxb6* and *Hoxc6*, as well as *Hoxa7* and *Hoxb7*, were ectopically expressed in the region anterior to the wing field. After 48 h, we observed either an anteriorly extended wing bud or a separated bud in the neck anterior to the endogenous wing bud (n = 226/440, Table 2). The efficiency of transfection and transcription was monitored by assessing EGFP expression from the plasmids used (Fig. 2K-O), and their wing-inducing effect by screening induced *Tbx5* expression (Fig. 2P-T). In more than half of the embryos, a separate wing bud, indicated by *Tbx5* expression, formed anteriorly to the endogenous wing bud (n = 128/226, Table 2). In the remaining embryos, the endogenous wing bud appeared extended anteriorly (n = 98/226, Table 2). These findings demonstrate that the ectopic formation of a wing bud in the neck is a consequence of the expression of all members of the HoxPG6/7 gene family.

**Table 2.** HoxPG6/7 upregulated wing bud formation in the neck region. Number indicate the numbers of embryos in which a cervical extension of the wing bud, or a cervical wing bud separated from the normal wing bud could be observed.

|  | A6 | B6 | C6 | A7 | B7 | Total |
| --- | --- | --- | --- | --- | --- | --- |
| Extension | 31 | 7 | 5 | 37 | 18 | 98 |
| Separated | 45 | 10 | 10 | 51 | 12 | 128 |
| Total | 76 | 17 | 15 | 88 | 30 | 226 |

Curiously, the induced wing buds did not grow distally to any great degree and remained small after 48h of re-incubation. To elucidate this phenomenon, RNA sequencing was used to compare gene expression in the induced wing buds with that of normal wing buds. Each group (Fig. 3A) was comprised of four replicates. Ectopic expression of *Hoxa6* resulted in the up-regulation of multiple genes shared with normal wing buds, and the gene expression pattern in *A6*-induced wing buds was more similar to that of cognate wing buds than to that of native neck tissue (Fig. 3B, B’). Gene Ontology (GO) biological process terms for 221 genes showed that the *A6*-induced bud closely resembles a normal wing bud (Fig. 3C, Table 3). Functional categorization revealed that 221 genes classified by GO biological process terms “anterior/posterior pattern specification” (p_genuine_ = 3.2^-10^; p_induced_ = 2.5^-10^), proximal/distal pattern formation” (p_genuine_ = 3.7^-9^; p_induced_ = 8.7^-9^), “regulation of transcription from RNA polymerase II promoter” (p_genuine_ = 8.4^-11^; p_induced_ = 3.2^-8^), “embryonic skeletal system morphogenesis” (p_genuine_ = 2.0^-9^; p_induced_ = 1.3^-5^), and “embryonic limb morphogenesis” (p_genuine_ = 8.1^-9^; p_induced_ = 1.4^-4^) were enriched in tissue of the genuine limb bud and in limb buds induced by *Hoxa6* overexpression (Table 4). In contrast, genes associated with the biological process terms “cell adhesion” (p = 4.3^-19^), “extracellular matrix organization” (p = 4.2^-15^), “transmembrane receptor protein tyrosine kinase signaling pathway” (p = 4.8^-13^), “positive regulation of kinase activity” (p = 1.6^-10^) and “multicellular organism development” (p = 3.1^-9^) were overrepresented among the genes enriched in neck tissue (Table 4). These findings demonstrate that *Hoxa6* is sufficient for wing bud induction.

**Table 3.**
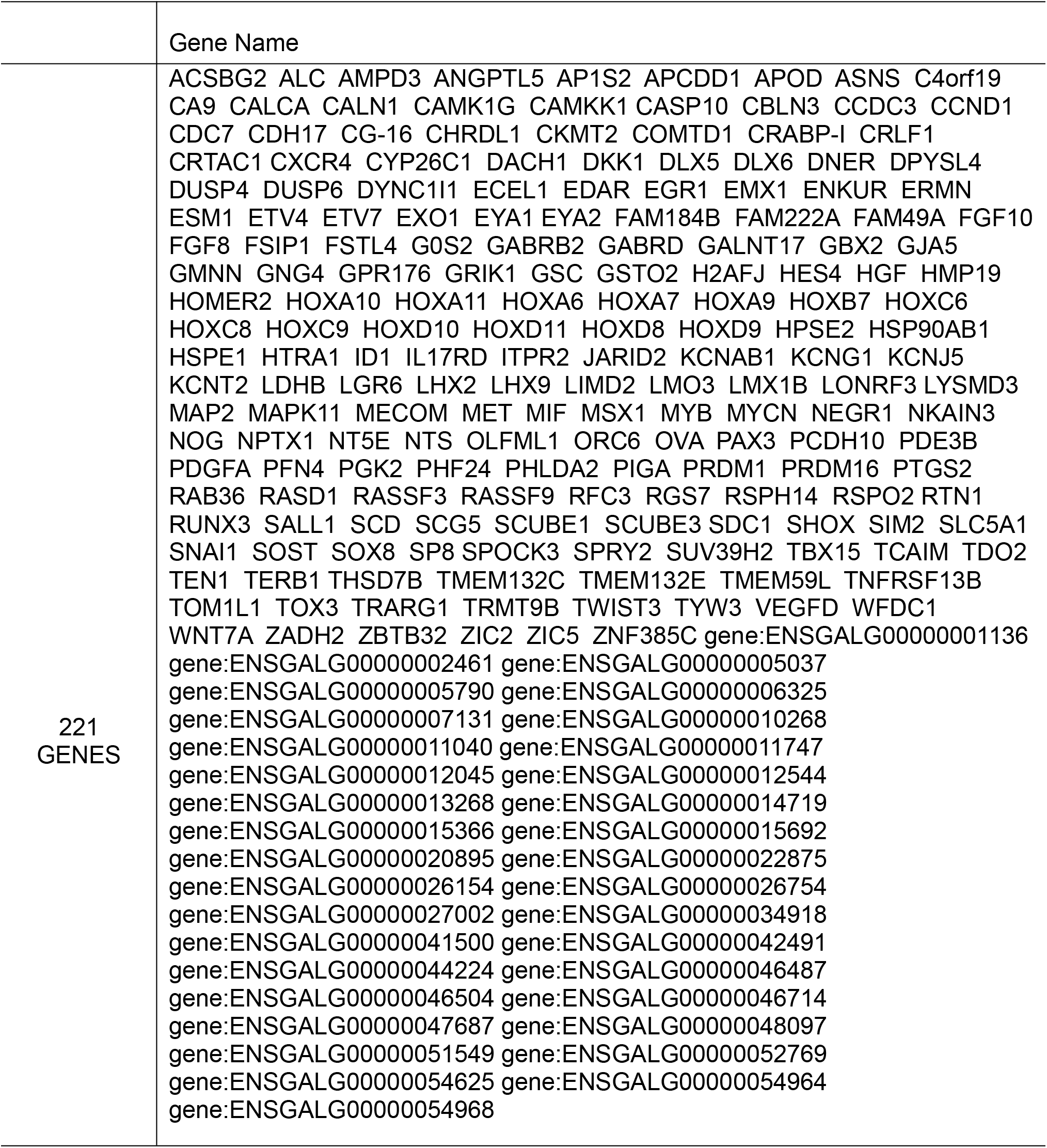
The name of 221 genes. 221 genes showed that the *A6*-induced bud closely resembles a normal wing bud.

**Table 4.** Gene Ontology analyses showing top ten terms in Biological Process.

| WING |  | A6-BUD |  | NECK |  |
| --- | --- | --- | --- | --- | --- |
| Term | p value | Term | p value | Term | p value |
| Regulation of transcription from RNA polymerase II promotor | 8.40E-11 | Anterior/posterior pattern specification | 2.50E-10 | Cell adhesion | 4.30E-19 |
| Anterior/posterior pattern specification | 3.20E-10 | Proximal/distal pattern formation | 8.70E-09 | Extracellular matrix organization | 4.20E-15 |
| Embryonic skeletal system morphogenesis | 2.00E-09 | Regulation of transcription from RNA polymerase II promotor | 3.20E-08 | Transmembrane receptor protein tyrosine kinase signalling pathway | 4.80E-13 |
| Proximal/distal pattern formation | 3.70E-09 | Protein folding | 1.50E-07 | Positive regulation of kinase activity | 1.60E-10 |
| Embryonic limb morphogenesis | 8.10E-09 | Embryonic skeletal system morphogenesis | 1.30E-07 | Multicellular organism development | 3.10E-09 |
| Dorsal/ventral pattern formation | 3.40E-08 | rRNA processing | 4.90E-05 | Cell-cell adhesion | 2.30E-08 |
| Neuron differentiation | 8.30E-08 | Embryonic limb morphogenesis | 1.40E-04 | Heart development | 4.20E-08 |
| Embryonic forelimb morphogenesis | 3.70E-06 | Ribosome biogenesis | 2.70E-04 | Axon guidance | 2.00E-07 |
| Embryonic hindlimb morphogenesis | 5.50E-06 | Neuron differentiation | 4.10E-04 | Negative regulation of cell migration | 1.20E-06 |
| Multicellular organism development | 8.60E-06 | Positive regulation of gene expression | 6.40E-04 | Blood coagulation | 1.40E-06 |

**Fig. 3.**
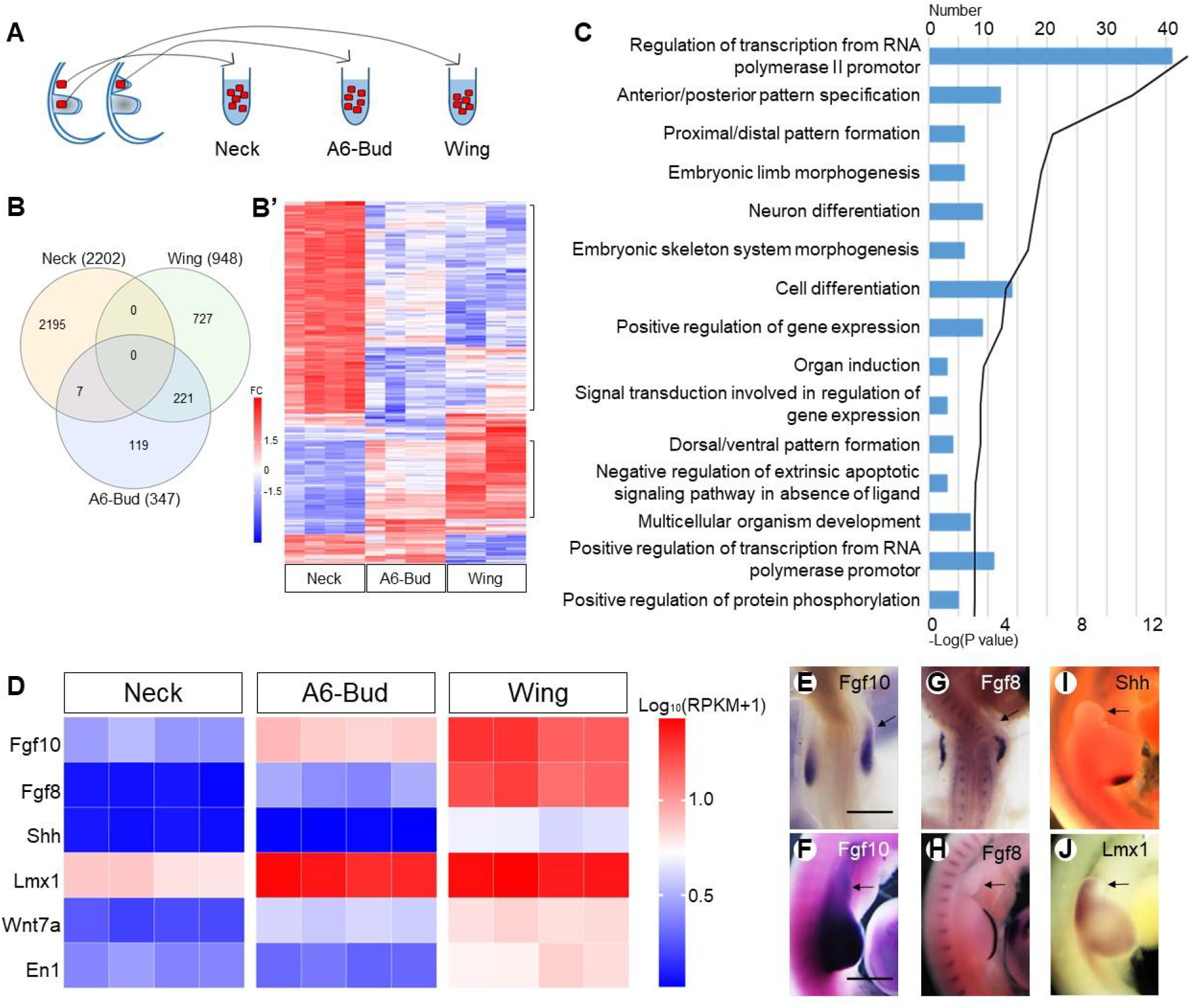
The neck wing bud is smaller than the natural wing bud. The scheme (**A**) indicates how tissue samples were collected for RNA sequencing. Venn diagram (**B**) showing the overlap between up-regulated genes expressed in normal wing bud (Wing 948), HoxA6-induced wing bud (A6-Bud 347) and neck tissue (Neck 2202) in the cervical LPM; the heatmap (**B’**) showing the expression profiles of genes in neck tissue, normal wing bud and HoxA6-induced wing bud. FC: fold change. Gene Ontology (**C**) analyses showing top 15 terms in biological process for 221 genes of A6-Bud. The heatmap (**D**) shown the expression levels of genes related to outgrowth, patterning. The expression of *Fgf10* (**E, F**), *Fgf8* (**G, H**), *Shh* (**I**) and *Lmx1* (**J**) in transfected embryos is rechecked by ISH. **E** and **G**: HH18-19; **F, H, I** and **J**: HH22; scale bars in **E** (for **E, G)** and in **F** (for **F, H**-**J)**: 500μm. Arrows indicate induced wing buds.

Although the wing program in *A6*-bud revealed by *Tbx5* was initiated, the AER was not established. Expression of *Fgf10* was activated in the neck, resulting in the initiation of mesodermal outgrowth. However, its expression level was lower than that of the physiological wing-forming mesoderm (Fig. 3D-F). In contrast, *Fgf8* was not induced in the ectoderm (Fig. 3D, G, H). Thus, the feedback loop between *Fgf10* and *Fgf8* was missing in the induced wing bud, and it failed to form an AER. Failure of the formation of functional AER is also indicated by the low levels of *Shh* expression in the induced wing bud as compared to the physiological wing anlage (Fernandez-Guerrero et al., 2022; Lin & Zhang, 2020). Without AER, the induced wing bud did not grow further. Further, the Zone of Polarizing Activity (ZPA) identified by the expression of *Shh* was not established (Fig. 3D, I). Finally, we noted that the induced wing bud was dorsalized, as indicated by the strongly upregulated expression of Lmx1 (Fig. 3D, J).

Taken together, we conclude that *HoxPG6/7* genes are sufficient for forelimb specification in the neck region. However, the induced wing bud is incapable of establishing the positive feedback loop between Fgf8 and Fgf10 due to the inability of Fgf signal transduction in the neck ectoderm (Lours & Dietrich, 2005).

## Discussion

In this study, we investigated how *Hox* genes determinate the forelimb cell fate of the LPM, thus the positioning of the forelimb. We found that functional inhibition of the A-cluster *Hox4/5/6/7* genes, on the protein level, using dominant-negative forms, resulted in reduction of *Tbx5* expression and subsequently of forelimb formation. Expression of *PG6/7* but not of *PG4/5 Hox* genes could reprogram neck mesoderm to limb-forming mesoderm. These findings indicate different roles of *PG6/7* and *PG4/5 Hox* genes during forelimb formation.

### *PG4/5/6/7* genes constitute the *Hox* code activating forelimb formation

In previous genetic studies, it has been shown that, in cooperation with Wnt and retinoic acid (RA) signalling (Nishimoto, Wilde, Wood, & Logan, 2015), *HoxPG4/5* genes activate *Tbx5* expression (Minguillon et al., 2012; Nishimoto et al., 2014; Moreau et al., 2019). *Tbx5* then activates *Fgf10* expression, which leads to the thickening and epithelio-mesenchymal transition of the LPM, initiating the formation of the primary forelimb bud (Delgado et al., 2021; Gros & Tabin, 2014). Subsequently, mesodermal *Fgf10* induces ectodermal *Fgf8* expression, creating a positive feedback loop that sustains the outgrowth of the limb bud. Experiments with dominant-negative forms suggest that not only *HoxPG4/5* but also *HoxPG6/7* are required for the *Tbx5* expression in the LPM and thus for forelimb formation. Functional inhibition of any of the A-cluster of *PG4/5/6/7 Hox* genes down-regulated *Tbx5*, as well as subsequent *Fgf10* and *Fgf8* expression. The resultant lower activity of the *Fgf10*-*Fgf8* feedback loop ultimately limited the further development of the wing buds.

In summary, our loss-of-function experiments provide direct evidences for the requirement of *PG4/5/6/7 Hox* genes for forelimb formation. Consequently, in addition to *PG4/5, PG6/7* genes also constitute the *Hox* code that activates the forelimb-forming program.

### *PG6/7* genes are sufficient for forelimb formation

Ectopic expression of *HoxPG6/7* genes activated *Tbx5* expression and initiated the wing-forming program in the neck LPM. Importantly, the induced wing bud in the neck did not grow sustainably. This may be linked to the reduced, or rather absent function of the FGF10-FGF8 feedback loop in the induced wing bud (Cohn & Tickle, 1999; Yin et al., 2016). The neck has previously been classified as a “limb-incompetent” region, where the limb formation can only occur when both limb mesoderm and limb ectoderm are simultaneously transplanted to the neck. Transplantation of limb mesoderm alone under neck ectoderm does not support limb formation (Lours & Dietrich, 2005). The re-specified wing mesoderm by *HoxPG6/7* in the neck is still covered by neck ectoderm. This condition is similar to the transplantation of the prospective limb mesoderm to the neck without limb ectoderm (Lours & Dietrich, 2005). Since the neck ectoderm is incapable of *Fgf* signal transduction, lacking Fgf8-Fgf10 feedback loop and AER, the development of the induced wing bud stalled in the pre-AER phase.

The wing buds seen following *PG6/7* expression in the neck resemble the wing anlagen in the chicken limbless mutant, in which *Fgf8* expression is mutated, and that lacks the AER and, like the induced limb buds here, the zone of polarizing activity (Grieshammer et al., 1996; Ros et al., 1996; Vogel et al., 1996). Moreover, both the induced neck wing-buds observed here and the wing buds of the limbless mutant are mainly dorsalized.

Importantly, implantation of FGF10-beads into neck LPM did not induce any wing bud structure in the neck (Lours & Dietrich, 2005). Neck wing buds can only be induced by ectopic expression of *HoxPG6/7* genes, as reported in the present study. This indicates that the emergence of ectopic limb buds from the neck requires re-specification of *Hox* code in the neck LPM. Despite the rudimentary outgrowth of the wing buds induced by ectopic *HoxPG6/7* expression in the neck region, our experiments demonstrate the pivotal role of *HoxPG6/7* in initiating the forelimb-forming program.

### *PG4/5* are insufficient for forelimb formation

Although both *PG4/5* and *PG6/7 Hox* genes impinge on *Tbx5* expression, they play different role during forelimb formation. In contrast to *HoxPG6/7*, neither the physiological expression of *HoxPG4/5* nor their overexpression in the neck region caused *Tbx5* expression and initiated formation of an ectopic wing bud. The distinct function of these two groups of *Hox* genes may be related to their expression pattern. The expression of *PG4/5* genes extends beyond the anterior border of the presumptive limb field and some of them are expressed in the entire neck region (http://geisha.arizona.edu). Accordingly, *Tbx5* is transiently activated in the entire neck region (Nishimoto et al., 2014). Yet this transient activation is inadequate to initiate forelimb formation, as normally no limbs originate from the neck region. It is only in the limb field where *PG4/5* expression overlaps with expression of *PG6/7* genes, that *Tbx5* expression is maintained and thus can initiate wing formation. Caudal to the forelimb region, this combinatorial effect is limited by *Hox9* expression (Cohn et al., 1997; Nishimoto & Logan, 2016; Tanaka, 2016). Functionally, *PG4/5 Hox* genes can activate *Tbx5* expression, but only the mesoderm expressing both *PG4/5* and *PG6/7 Hox* genes can form forelimb. Similar findings have been observed in the specification of motor neurons for the forelimb skeletal muscles (Mukaigasa et al., 2017). The early forelimb motor neuron programme starts in the entire neck region, but only motor neurons under the control of *Hox4/5* and *Hoxc6* complete their differentiation. Neurons solely under the control of *Hox4/5* undergo apoptosis.

### Redundancy of limb-forming *Hox* genes

The partial reduction of wing development seen after dominant-negative form expression with downstream action of any of the *Hoxa4/5/6/7* genes is fully consistent with the partial redundancy among *Hox* paralog groups described for *HoxPG5* and *HoxPG6* during axial patterning (McIntyre et al., 2007) and *HoxPG5* in limb development (Xu et al., 2013). We note, though that we cannot formally exclude incomplete blockade of the genes targeted given the competitive nature of our approach. Be that as it may, our *Hox*-inactivation experiments clearly reveal a dosage effect of *Hox* genes on orthologous limb development. They further lead to the conclusion that normal wing development may depend on the balanced expression of *HoxPG4/5/6/7* genes.

The absence of overt limb phenotypes in PG4–PG7 mouse mutants likely reflects both the extensive functional redundancy among Hox paralogs and the difficulty of detecting subtle limb-specific effects in bilateral, systemically affected embryos. In contrast, the chick embryo system allows unilateral gene manipulation, providing an internal control and greater sensitivity for detecting weak or localized effects that may be masked in whole-animal mouse mutants. This difference in experimental sensitivity likely explains why limb phenotype that are undetectable in mouse mutants can be clearly revealed by targeted manipulations in the chick model.

### Permissive and instructive mechanisms during limb evolution

It has been hypothesized that an interplay between permissive, instructive and inhibitory mechanisms is needed to induce precise tissue organization (Morales et al., 2021). Such an interplay may also regulate limb positioning. As shown by several authors, the caudal boundary of the forelimb is determined through the antagonism of the rostral and caudal codes: the rostral code induces forelimb formation, whereas the caudal code inhibits it (Cohn et al., 1997; Moreau et al., 2019; Nishimoto & Logan, 2016; Nishimoto et al., 2014; Tanaka, 2016). In the present study, we suggest that the rostral code should comprise two functionally distinct subgroups. Our data show that inhibiting HoxPG4/5 disrupts limb formation, indicating its necessity. However, overexpressing *HoxPG4/5* alone does not induce limb formation, suggesting they are not sufficient. In contrast, *HoxPG6/7* is both necessary and sufficient, as their inhibition prevents limb formation and their overexpression induces limb formation.

Thus, we speculate that *HoxPG4/5* set up a permissive environment by initiating transient *Tbx5* expression that allows limb formation to occur but does not directly trigger the process. The broad expression domain of *HoxPG4/5*, including the neck region, defines an extended permissive region where forelimb formation might be initiated. In contrast, *HoxPG6/7* instructively directs the formation of limbs by maintaining *Tbx5* expression in a precise position. The overlap of *HoxPG6/7* expression domains with the limb field further supports instructive roles of these *Hox* genes.

Moreover, the extended *Tbx5* expression domain from the heart to forelimb signifies the posterior shift of the forelimb (Anderson et al., 2016) (Fig. 4). As the forelimb programme proceeds, *Tbx5* expression is maintained in only the heart and the prospective forelimb region (Fig. 4). Notably, the regression of *Tbx5* in the neck region between the heart and forelimb region implies functional differences between *PG4/5* and *PG6/7* genes.

**Fig. 4.**
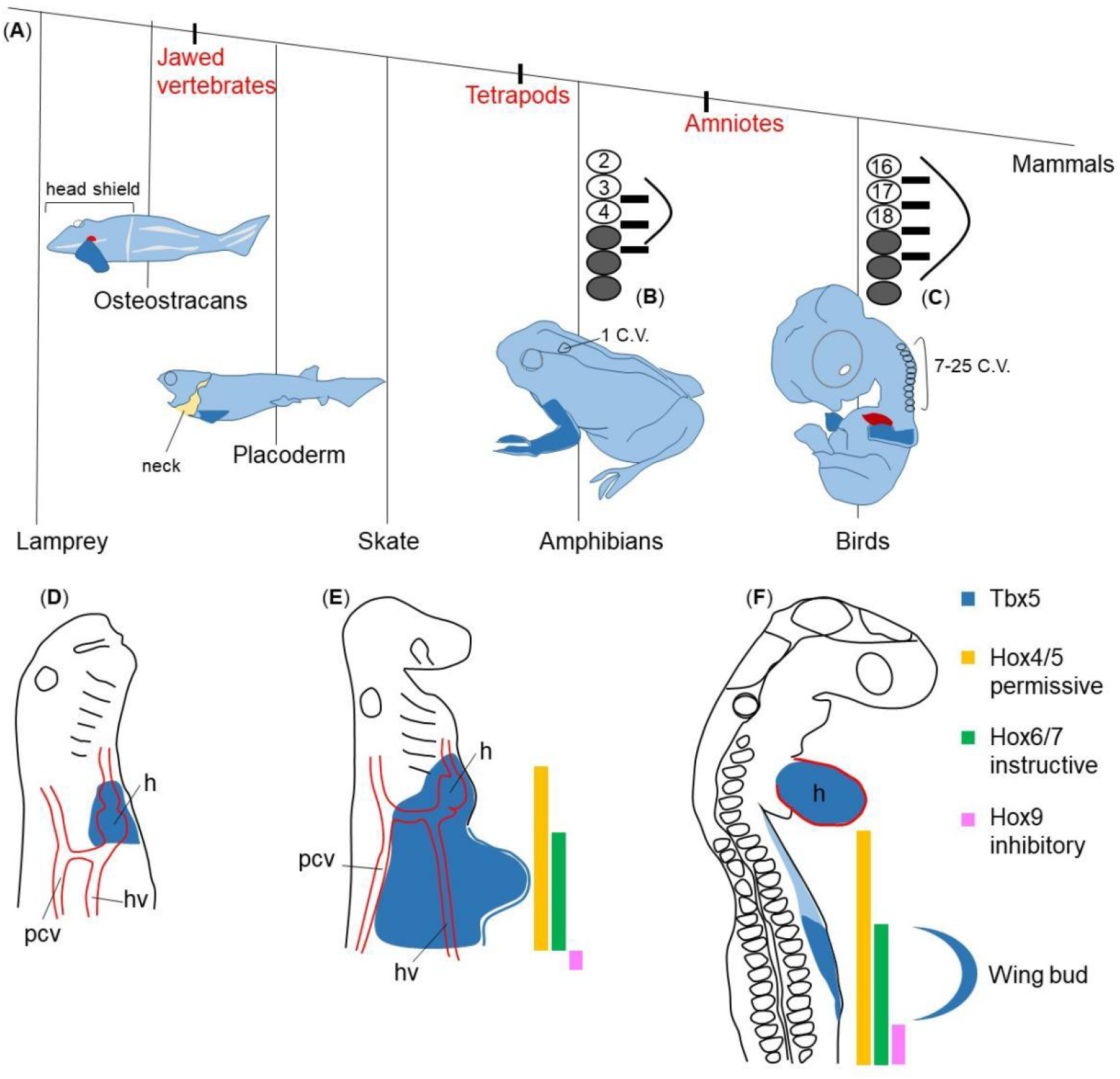
Permissive, instructive, and inhibitory *Hox* codes regulate the forelimb positioning. The phylogenetic tree of gnathostomes (**A**, redrawn from Hirasawa et al., 2016) show that despite the variation in the number of cervical vertebrae (C.V.), the pectoral fin and forelimb (dark blue) are always located at the cervical–thoracic boundary. However, their axial positions with respect to somite number vary widely across species (**B** and **C** modified from Burke, 1995). **B, C**: Bright circles: numbered somites; grey shaded circles: thoracic somites; black bars: spinal nerves of the brachial plexus; curved lines: limb bud. In lamprey embryos **(D)**, expression of *Tbx5* homologue is restricted to the heart region (Adachi et al., 2016). In skate embryos (**E**), *Tbx5* expression (blue) extended slightly caudally from the heart anlage (Adachi et al. 2016). In avian embryos (**F**), it extends from the heart over the neck to the wing field. During wing bud formation, *Tbx5* expression is restricted to only the heart and the wing bud. In lateral plate mesoderm, *Hox4/5* expression (yellow) extends into the neck region, whereas the anterior expression domain of *Hox6/7* (green) is at the wing level. *Hox9* expression (magenta) starts posteriorly to the wing. The yellow, green, and magenta colours represent permissive, instructive, and inhibitory functions, respectively. The blue curved line outlines the wing bud.

There is an evolutionarily conserved requirement for spatial and temporal regulation of cell behaviour during morphogenesis. *Hox* codes control the growth and shape of almost all organs and the body as a whole. Therefore, the identified mechanisms by which the Hox code genes play permissive and instructive roles in controlling cell behaviour are of general significance for organogenesis during embryonic development and adult regeneration and may elucidate the regional specification mechanisms for other organs.

### Towards an evolutionary perspective of vertebral morphology and limb positioning

While the length of the cervical spinal column and the position of the forelimbs are highly fixed in mammals, they are much more variable in other vertebrates, especially from an evolutionary perspective. Indeed, there appears to be an evolutionary trend toward increased head mobility, achieved through the increasing complexity and length of the cervical spine. This trend involves, or presupposes, a caudal repositioning of the anterior limbs.

Extant jawless vertebrates such as lampreys and hagfish lack any morphological vestige suggesting an anlage of anterior limbs. In these species, the expression of Tbx4/5, the hallmark marker of incipient anterior limb and heart development, is restricted to the latter (Adachi, 2016) (Fig. 4D). The first pectoral fins, defined by the presence of a possibly gill-arch-derived pectoral girdle (Janvier, 1996) and connected to the head shield, are found in fossil osteostracans, an early class of gnathostomes (Coates, 1994) (Fig. 4A).

The separation of the pectoral girdle from the head shield resulted in the development of a primary neck, first identifiable in placoderms (Trinajstic et al., 2013) (Fig. 4A). In jawed fish with paired fins, the evolutionary caudal repositioning of the anterior pectoral fins can also be verified by the fact that the expression of Tbx5 is now slightly caudal to the heart anlage (Anderson et al., 2016). This has been documented in skates as well as zebrafish (Adachi et al., 2016; Criswell et al., 2021) (Fig. 4E).

A true neck connecting the cranium and trunk first evolved in amphibians—the first land vertebrates—as the pectoral girdle shifted caudally and the first trunk vertebra transformed into a cervical vertebra (Torrey, 1978) (Fig. 4A, B). With the further evolution of land vertebrates, the number of cervical vertebrae increased significantly (Goodrich, 1906). The longest cervical vertebral columns, with 76 segments, have been reported in the fossil diapsids Muraenosaurus and Elasmosaur Albertonectes (Kubo et al., 2012; Young, 1981). In birds, the number of cervical vertebrae varies widely, ranging from nine to twenty-five (Yapp & Lyons, 1965) (Fig. 4A, C, E). The evolutionarily retained muscular connection between the head and shoulder girdle, formed by the cucullaris muscle and its derivatives, validates this history (Sefton et al., 2016; Theis et al., 2010).

The significance of Hox genes in vertebrate diversification and limb complexity has been repeatedly documented (Cohn & Tickle, 1999; Wellik & Capecchi, 2003; Li et al., 2023; Korth & Polly, 2023). The present results refine our understanding of how Hox genes integrate vertebral column structure and limb positioning, which together have led to the extensive behavior and foraging/predatory diversification of vertebrates (Rytel et al., 2024; Marek et al., 2021).

## Materials and Methods

### *In ovo* electroporation

Fertilised chicken (Gallus gallus domesticus) eggs were obtained from the Institute of Animal Sciences of the Agricultural Faculty, University of Bonn, Germany. First, after windowing of the egg shell and exposing the embryo, a solution containing 5–10 µg/µL plasmid and 0.1 % Fast Green was injected into the coelom at specific axial levels. Electroporation was then performed using the CUY 21-Edit-II electroporator with one poration pulse of high voltage (0.01 ms, 70 V) followed by two driving pulses of low voltage (50 ms, 7 V, with 200 ms intervals). There is a 99.9 ms interval between the high and low voltage pulses. After reincubation, embryos were imaged under the Nikon SM21500 fluorescence microscope and then fixed in 4 % paraformaldehyde overnight at 4°C.

### Plasmids for electroporation

DNA plasmids were produced by Dongze Bio-products (Guangzhou, China). Coding sequences (obtained from NCBI) for Hoxa4 (930bp, NM_001030346.3), Hoxa5 (813bp, NM_001318419.2), Hoxa6 (696bp, NM_001030987.4), Hoxb6 (669bp, NM_001396636.1), Hoxc6 (714bp, NM_001407494.1), Hoxa7 (660bp, NM_204595.3) or Hoxb7 (654bp, XM_040653307.2) were inserted into the pCAGGS-P2A-EGFP plasmid. A plasmid expressing the dominant negative (dn) form specific for Hoxa4, a5, a6 or a7 was produced using their coding sequence lacking the C-terminal portion, including Hoxa4dn (762bp), Hoxa5dn (729bp), Hoxa6dn (585bp) and Hoxa7dn (528bp). A large quantity of DNA plasmids was purified using the NucleoBond Xtra Midi DNA preparation kit (MACHEREY-NAGEL).

### RNA in situ hybridisation

Whole-mount RNA in situ hybridization was performed by incubating probes at 65°C (Nieto, Patel et al. 1996). The probes were detected using anti-Digoxigenin-AP, fab fragments (Roche) and color reagent NBT/BCIP staining solution (Roche). Chicken Lmx-1, Fgf10 and Fgf8 probes were provided by H. Ohuchi, O. Pourquie and C. Tabin, respectively. Chicken Tbx5 probe, Hox probes and Hoxdn C-terminal probes were produced using PCR and transcribed using the DIG-RNA Labelling Kit (Roche, #11175025910) with T7 polymerase. The specific primers were shown in Table 5.

**Table 5.**
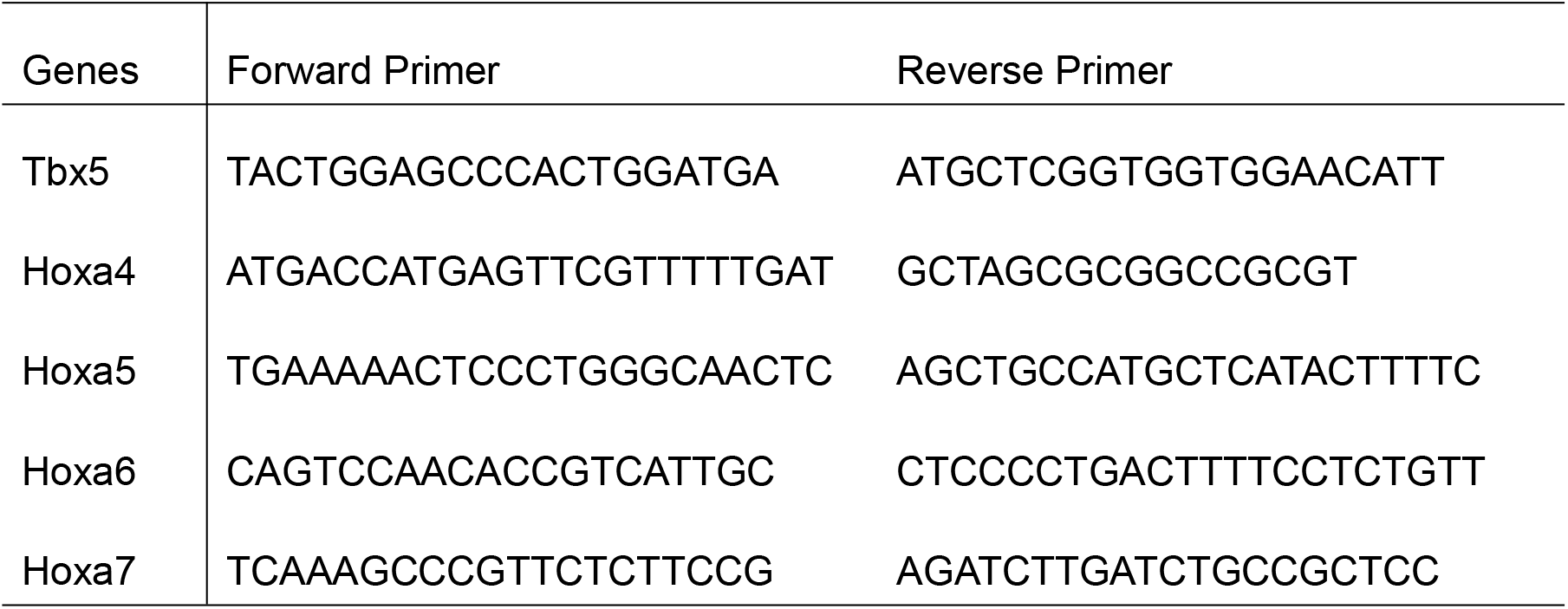
Primer sequences used for generating in situ hybridization probes by PCR.

### RNA-Seq analyses

Wing parts (five samples per replicate, four replicates, total 20 samples) and neck parts (20 samples per replicate, four replicates, total 80 samples) were dissected from HH22 normal embryos. Additionally, a total of 80 *Hoxa6*-induced ectopic buds (20 samples per replicate, four replicates) were dissected from HH22 embryos with *Hoxa6* ectopic expression in the neck. The dissections were performed under the Nikon SM21500 fluorescence microscope. Only ectopic buds identified by their morphology and EGFP expression were isolated and collected, including the surface ectoderm. Total RNA of samples was isolated with miRNeasy Micro Kit (QIAGEN). Library preparation was performed according to the manufacturer’s protocol using the ‘VAHT Universal RNA-Seq Library Prep Kit for Illumina V6 with mRNA capture module’. Next, 500 ng total RNA was used for mRNA capturing, fragmentation, cDNA synthesis, adapter ligation and library amplification. Bead-purified libraries were normalised and finally sequenced on the HiSeq 3000/4000 system (Illumina Inc. San Diego, USA).

### Statistical analysis

Data analyses on FASTQ files were conducted with CLC Genomics Workbench (version 21.0.4, QIAGEN, Venlo. NL). The reads of all probes were adapter trimmed (Illumina TruSeq) and quality trimmed. Mapping was done against the Gallus gallus (GRCg6a) (19 March, 2021) genome sequence. Statistically significant differential expression was determined using the ‘Differential Expression for RNA-Seq’ tool (version 2.4) (Qiagen Inc. 2021). The resulting *P* values were corrected for multiple testing by FDR. The RNA expression level was indicated by reads per kilobase of transcript per million mapped reads (RPKM) and the statistical analysis between the three groups were made by ordinary one-way ANOVA, using GraphPad Prism v6 (San Diego, CA, USA). Functional annotation clustering was done by means of the DAVID online tool (https://david.ncifcrf.gov/) and using the Gene Ontology “biological process” annotation category. Data are presented as mean ± standard error of the mean. The level of statistical significance was set at **p < 0.01.

## Acknowledgments

The authors thank Dr. Frank Stockdale for helpful discussions and valuable comments on the manuscript. We thank Sandra Graefe and Heinz Bioernsen for their expert technical assistance. Computational support from the Centre for Information and Media Technology, especially the High-Performance Computing team at Heinrich-Heine University, is acknowledged. This work was supported by grants from China Scholarship Council (CSC) and by German Research Funding (DFG-Hu 729/13).

## Author Contributions

Conceptualisation: YW, RH, KP, MH, TB, SK, QP. Investigation: YW. Methodology: YW, JBW, LC, JLW, PZ, HT, ZL, XQ, DC. RNA-Seq: PP, KK, YW. Funding acquisition: YW, SK, RH. Writing: YW, RH, KS, MH, KP, SK, QP, JW, GK.

